# Self-organizing glycolytic waves fuel cell migration and cancer progression

**DOI:** 10.1101/2024.01.28.577603

**Authors:** Huiwang Zhan, Dhiman Sankar Pal, Jane Borleis, Chris Janetopoulos, Chuan-Hsiang Huang, Peter N. Devreotes

**Author notes:** Corresponding authors: Huiwang Zhan, Chris Janetopoulos, Chuan-Hsiang Huang, and Peter N. Devreotes.

## Abstract

Glycolysis has traditionally been thought to take place in the cytosol but we observed the enrichment of glycolytic enzymes in propagating waves of the cell cortex in human epithelial cells. These waves reflect excitable Ras/PI3K signal transduction and F-actin/actomyosin networks that drive cellular protrusions, suggesting that localized glycolysis at the cortex provides ATP for cell morphological events such as migration, phagocytosis, and cytokinesis. Perturbations that altered cortical waves caused corresponding changes in enzyme localization and ATP production whereas synthetic recruitment of glycolytic enzymes to the cell cortex enhanced cell spreading and motility. Interestingly, the cortical waves and ATP levels were positively correlated with the metastatic potential of cancer cells. The coordinated signal transduction, cytoskeletal, and glycolytic waves in cancer cells may explain their increased motility and their greater reliance on glycolysis, often referred to as the Warburg effect.

## Introduction

Classical biochemistry holds that enzymes of the glycolytic pathway reside in the cytosol and that glycolysis takes place within the cytoplasm. Consistently, the enzymes separate into the supernatant of fractionated cells; indeed the release of soluble aldolase is often considered as a marker for cell lysis ^1^. Furthermore, glycolytic activity, as measured by the conversion of glucose to pyruvate, can be reconstituted *in vitro* by mixing purified enzymes and substrates in appropriate concentrations in solution ^2^. In cells, pyruvate enters mitochondria from the cytosol and initiates oxidative phosphorylation. Although oxidative phosphorylation yields more ATP than glycolysis, the relative contribution of the two processes toward ATP production varies among cell types. It was found in the 1920s that cancer cells often shift to a greater dependence on glycolysis, a phenomenon referred to as the Warburg effect ^3–5^.

Recent studies have suggested alternate localizations or configurations of some glycolytic enzymes. Aldolase is known to be associated with F-actin in vitro ^6–9^. Aldolase was reported to be released from stress fibers by PI3 Kinase and the small GTPase Rac ^10^. In another study, phosphofructokinase (PFK) level was found to be regulated by tension through E3 ubiquitin ligase associated with stress fibers ^11^. Glycolytic activity has also been reported to localize to synapses during stress and neuronal growth cones to provide energy for their induced collapse ^12,13^ as well as the invasive protrusions of anchor cells in *Caenorhabditis elegans* ^14^. Additionally, PFK can, under certain conditions, polymerize into short fibrils and that this configuration was important for cell migration ^15^. These studies suggest that the localization of glycolytic enzymes in cells may not be exclusively cytosolic as inferred by the early biochemical experiments on broken cells. Dynamic localization may play an important regulatory role in addition to various allosteric regulators that have been described.

Energy intensive cellular activities such as phagocytosis, cytokinesis, and cell migration require cell shape changes induced by signal transduction and cytoskeletal activities, which bring about various membrane protrusions and deformations. Emerging studies in many different types of cells have unveiled fascinating phenomena underlying these protrusive events. Waves of signal transduction and cytoskeletal activity propagate across the cell membrane and cortex matching closely the protrusive events ^16–24^. Perturbation of the wave behavior has a predictable, causal effect on the protrusions and dependent physiological events ^25–27^. The propagating waves are manifestations of biochemically excitable networks involving Ras/PI3K signaling and anionic phospholipids which drive dynamic changes in actin and actomyosin networks ^28–30^. Interestingly, there is an increased frequency of these waves in more aggressively metastatic cells ^26,31^. The increased dependence on glycolysis and the higher frequency of cortical waves in cancer cells raises intriguing questions about the possible role of the waves in the regulation of glycolytic enzymes and the implications for cellular function.

These observations prompted us to further investigate the relationship between the waves and the glycolytic enzymes in normal human and metastatic mammary epithelial cells. Surprisingly, we found that most of the glycolytic enzymes were associated with the cell cortex, and co-localized with the cortical waves. Increasing wave activity resulted in enzyme recruitment while abolishing the waves returned the enzymes to the cytosol. Consistently, ATP levels were strongly correlated with the augmentation or abrogation of the waves. Remarkably, recruitment of a single glycolytic enzyme induced epithelial cell spreading and, in neutrophils, accelerated migration, while the inhibiting of glycolysis silenced cellular protrusions. Furthermore, recruitment of PFK causes a co-recruitment of aldolase, suggesting that enzymes might be linked to a complex. Notably, there was an increased frequency of waves and a concomitant increased level of ATP in more metastatic cells, indicating that the higher abundance of waves of glycolytic activity observed in cancer cells may explain the elevated metabolic activity associated with the Warburg effect.

## Results

### Glycolytic enzymes are enriched at Lifeact labeled waves and protrusions

To visualize the localization of glycolytic enzymes, we first introduced GFP-tagged aldolase into the mammary epithelial cell MCF-10A M3. We captured images of the cells’ basal surfaces and made a fascinating observation: while the aldolase was distributed within the cytosol as expected, a significant fraction was found to be associated with dynamic waves moving across the basal surface of the cell (**Figures 1A and S1A, Video S1**). Line kymographs drawn through several different planes exhibited the dynamic spatial and temporal characteristics of aldolase localization in the propagating waves (**Figures 1B and S1B**).

**Figure 1.**
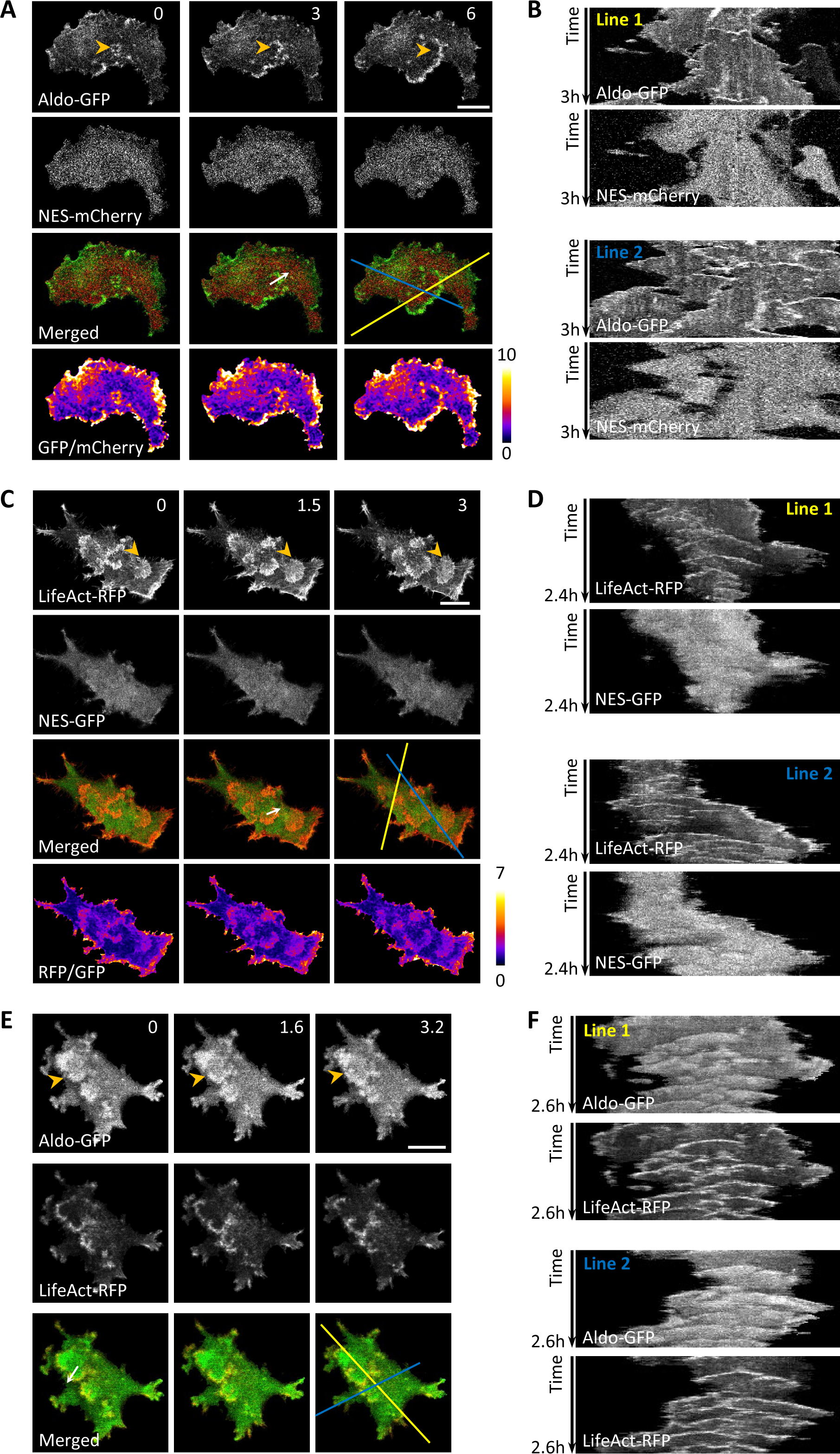
Enrichment of aldolase in F-actin waves and protrusions. (A) Confocal images of the basal surface of an MCF-10A M3 cell expressing aldolase-GFP and NES-mCherry (also see **Video S1**). Another cell example is shown in **Figure S1A**. (B) Kymographs of fluorescence intensity along the two lines in (A) over time. (C) Confocal images of the basal surface of a cell expressing LifeAct-RFP and NES-GFP (also see **Video S2**). (D) Kymographs along the two lines in (C) over time. (E) Confocal images of the basal surface of a cell expressing aldolase-GFP and LifeAct-RFP (also see Video S3). (F) Kymographs along the two lines in (E) over time. The orange arrowheads in (A), (C), and (E) indicate expanding waves propagating across the basal surface of the cell. The scan of fluorescence intensity across the white arrows in (A), (C), and (E) are shown **Figures S1D, E,** and **F**, respectively. Scales bars are 20 μm and the unit of time stamp is min throughout all figures unless otherwise indicated.

In the same cells, mCherry was nearly evenly spread throughout the cytosol. Merged images revealed a clear enrichment of aldolase in the waves compared to the cytosolic mCherry signal. To quantitatively assess the enrichment in the waves, we normalized the aldolase-GFP signal to the cytosolic mCherry signal, revealing a significantly heightened signal within the aldolase waves (**Figure 1A**, **bottom**). Scans across the wave area of the cell showed there was no increase in the cytosolic mCherry control within the waves, but a large increase for aldolase, with a 2 to 10-fold higher ratio compared to other cytosolic regions (**Figure S1C** **and S1D**).

We performed the analogous analyses for the distribution of Lifeact, a marker for F-actin, which is also localized to propagating waves on the basal surface of cells (**Figures 1C**, **Video S2**). Merged and ratio images by normalizing the Lifeact signal to cytosolic GFP further supported the enhancement of Lifeact in the wave region. Line kymographs also showed the dynamic nature of the F-actin waves, absent in the GFP control (**Figure 1D**). Line scans indicated a 3-fold increase of Lifeact in the waves (**Figure S1E**).

Interestingly, when aldolase-GFP and Lifeact-RFP were co-expressed, both markers were enriched in traveling waves and protrusions that emerged when a wave reached the cell’s edge (**Figure 1E, Video S3**). Examination of merged images revealed that the distribution of aldolase was consistently more diffuse compared to that of Lifeact (**Figure 1E**). Line kymographs illustrated similar spatial-temporal patterns of aldolase and Lifeact in the propagating waves and protrusions (**Figure 1F**). Line scans indicated that the half-width of the aldolase waves was approximately twice that of the F-actin waves (**Figure S1F**).

The presence of aldolase in waves prompted further investigation into the cellular localization of other glycolytic enzymes (**Figure 2A**). Of the eight remaining enzymes in the glycolysis cascade, five, namely hexokinase (HK), phosphofructokinase (PFK), glyceraldehyde 3-phosphate dehydrogenase (GAPDH), enolase, and pyruvate kinase (PK), could be tagged with GFP or RFP for visualization and successfully expressed in the epithelial cells. All five glycolytic enzymes tested exhibited enrichment within actin waves and protrusions (**Figures 2B-K, Videos S4-8**). Co-expressing each glycolytic enzyme with LifeAct, labeled with an appropriate complementary color, enabled simultaneous imaging, highlighting the dynamic localization patterns in waves and protrusions. Phosphofructokinase (PFK) demonstrated the strongest enrichment within the waves, similar to aldolase. Some enzymes, such as GAPDH, appeared to be more diffuse than PFK but, nevertheless, were clearly enriched above the cytosolic level. The temporal and spatial patterns of glycolytic enzyme localization showed high coordination with waves of actin polymerization, evident from color-coded temporal overlay images (**Figures 2C, E, G, I, K**). Scans through the waves showed that, as found for aldolase, the other glycolytic enzymes display more diffusive patterns than LifeAct (**Figure 2L**).

**Figure 2.**
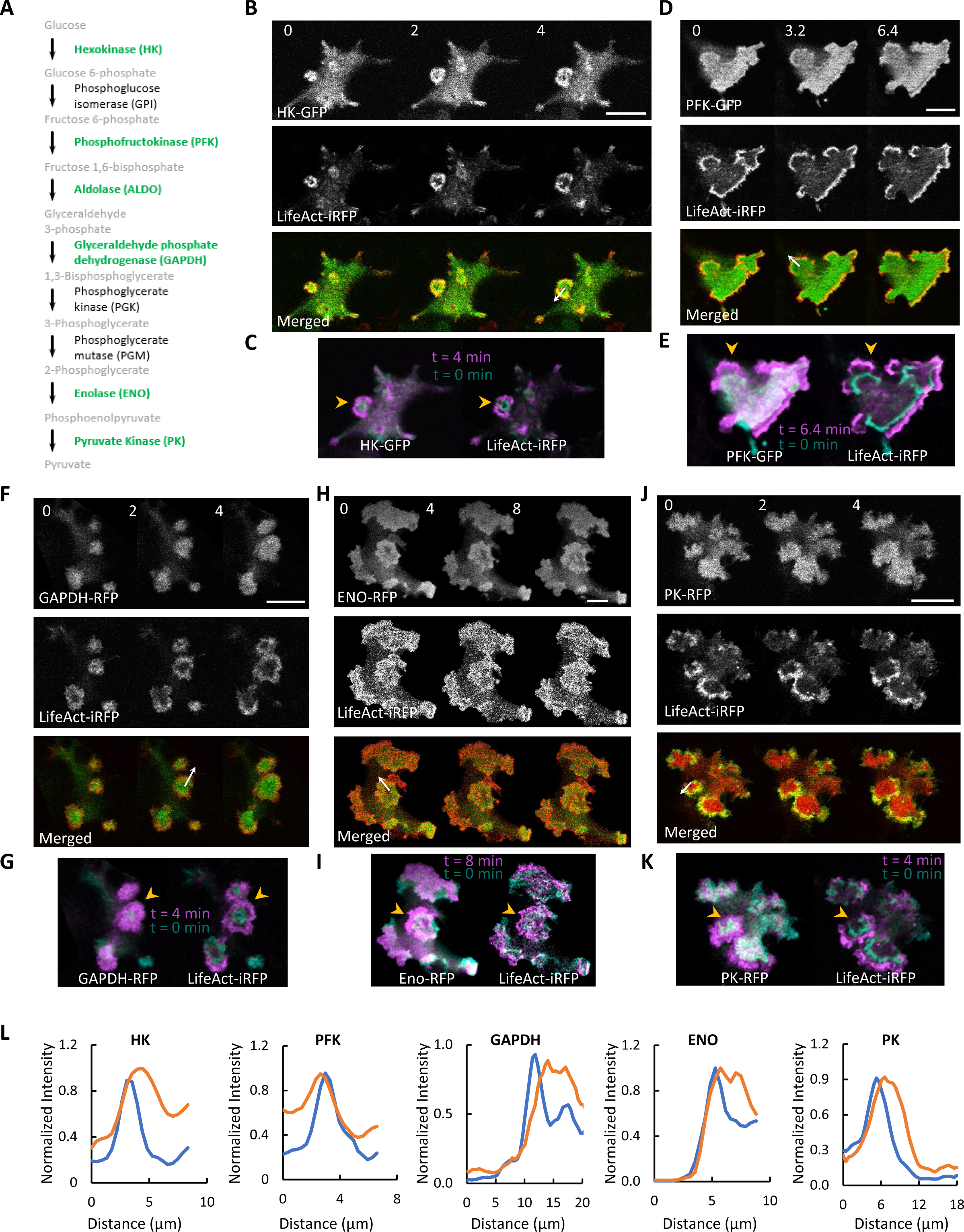
Enrichment of additional glycolytic enzymes in F-actin waves and protrusions. (A) The glycolytic pathway. Enzymes that could be fluorescently tagged and imaged in this study are shown in green. (B-K) Colocalization of LifeAct and glycolytic enzymes in waves. Confocal images of the basal surface of MCF-10A M3 cells expressing LifeAct-iRFP670 and HK-GFP (B), PFK-GFP (D), GAPDH-RFP (F), enolase-RFP (H), and PK-RFP (J) are shown (also see **Videos S4-8**). Color-coded overlays show the progression of waves over time (C, E, G, I, K). The orange arrowheads in (C), (E), (G), (I) and (K) indicate expanding waves propagating across the basal surface of the cell. Full length proteins are used except for HK, in which the first 21 a.a. of the N-terminus were truncated ^55^. (L) Normalized intensity of LifeAct-iRFP (blue) and FP-tagged glycolytic enzymes (orange) across the white arrow in (B, D, F, H, J).

While not all glycolytic enzymes were expressed in this study, our results suggest a trend of enrichment for the entire glycolysis cascade within actin waves that generate ruffles in cells ^17,18,26,32^. The dynamic clustering of enzymes within these waves is anticipated to significantly elevate the concentration, thereby substantially augmenting the glycolysis rate. This observation implies that glycolytic enzymes could play an active role in actin-based structural formations, potentially impacting the generation of cellular protrusions.

### Altering waves of glycolytic enzymes causes parallel changes in glycolytic activities

To further investigate the correlation between the glycolytic enzymes and cortical waves, we examined the activities of glycolytic waves under perturbations that are known to change the wave dynamics. It was previously shown that addition of EGF and insulin increases the Ras/PI3K and actin wave activity in these cells within minutes and this effect persists for hours ^26^. We monitored GFP tagged aldolase and PFK in the ventral surface of the MCF-10A M3 cells before and after stimulation with EGF and Insulin. While aldolase waves were present prior to the addition of EGF, the number of waves dramatically increased with the addition of EGF and Insulin (**Figure 3A** **and Video S9**). The response was variable among individual cells (**Figure 3B**), but on average there was a ∼2-fold increase in activities which plateaued within 30 min and was maintained for the time imaged (**Figure 3C**). Similar effects were observed for PFK (**Figures 3D-F** **and Video S9**).

**Figure 3.**
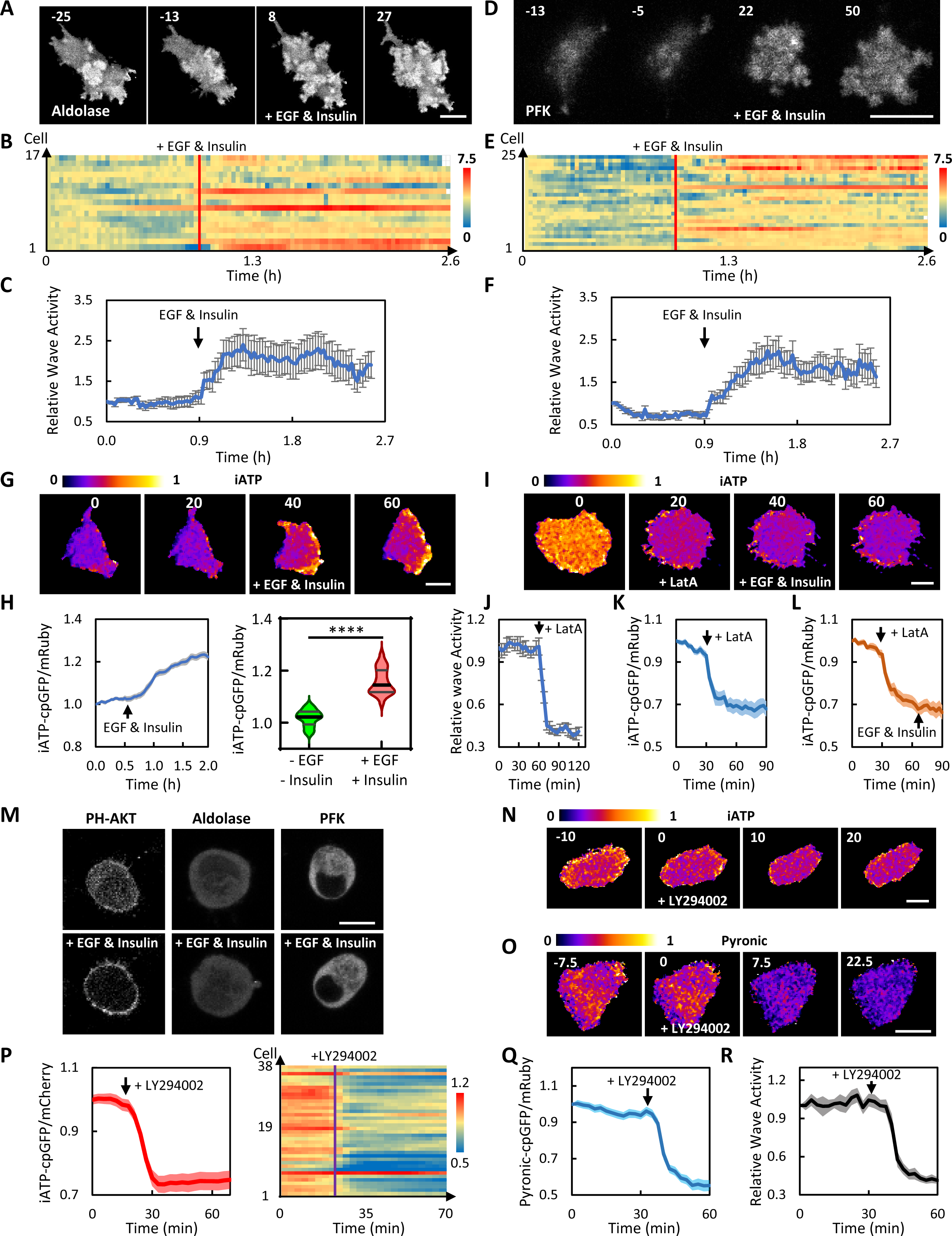
Effects of perturbing glycolytic waves. (A) Confocal images of the basal surface of a cell expressing aldolase-GFP stimulated with 20 ng/ml EGF and 10 μg/ml insulin at 0 min (also see **Video S9**). (B) Wave activity, defined as areas of pixels with above threshold aldolase-GFP intensity, of cells stimulated with EGF and insulin. The wave activity normalized to that of the first frame was plotted over 2.6 h for 17 individual cells. (C) Plot of normalized wave activity (mean ± SEM) over time for the 17 cells in (B). (D) Confocal images of the basal surface of a cell expressing PFK-GFP stimulated with 20 ng/ml EGF and 10 μg/ml insulin at 0 min (also see **Video S9**). (E) PFK-GFP wave activity of cells stimulated with EGF and insulin. The wave activity normalized to that of the first frame was plotted over 2.6 h for 25 individual cells. (F) Plot of normalized wave activity (mean ± SEM) over time for the 25 cells in (E). (G) iATP cpGFP/mRuby ratio images of a cell stimulated with EGF and insulin. (H) Left panel: Plot of normalized iATP cpGFP/mRuby (mean ± SEM) over time for 21 cells stimulated with EGF and insulin. The dynamic changes in iATP cpGFP/mRuby of these 21 individual cells were plotted in **Figure S4A**. Right Panel: Violin plots of average iATP cpGFP/mRuby ratios before and after EGF and insulin stimulation in these 21 cells. ****p <0.0001 (paired t test). (I) iATP cpGFP/mRuby ratio images of a cell treated with 10 μM Latrunculin A (LatA) followed by stimulation with EGF and insulin. (J) Plot of normalized ratio of membrane to cytosol LifeAct (mean ± SEM) over time for 15 cells treated with LatA. (K) Plot of normalized iATP cpGFP/mRuby (mean ± SEM) over time for 20 cells treated with LatA. The activity of individual cells was plotted in **Figure S4B**. (L) Plot of normalized iATP cpGFP/mRuby (mean ± SEM) over time for 23 cells treated with LatA followed by EGF and insulin. The activity of individual cells was plotted in **Figure S4C**. (M) Confocal images of PH-AKT-RFP, aldolase-GFP, and PFK-GFP before and after treatment with EGF and insulin in cells pretreated with 10 μM LatA. (N) iATP cpGFP/mRuby ratio images of a cell treated with 50 μM LY294002 at 0 min. (O) Pyronic cpGFP/mRuby ratio images of a cell treated with 50 μM LY294002 at 0 min. Images of other channels of this cell are shown in **Figure S4E**. Also see **Video S11**. (P) Plot of normalized iATP cpGFP/mRuby (mean ± SEM) over time for 38 cells treated with LY294002. Dynamic changes of individual cells are plotted in the right panel. (Q) Plot of normalized pyronic cpGFP/mRuby (mean ± SEM) over time for 23 cells treated with LY294002. Activities of individual cells are shown in **Figure S4F.** (R) Plot of normalized ratio of membrane to cytosol LifeAct (mean ± SEM) over time for 16 cells treated with LY294002. Cells showing changes in iATP biosensor upon treatment with EGF and Insulin, LatA, and LY 294002 were all pretreated with OAR. Scale bar is 20 μm for all except (M) (10μm). All the cells plotted and quantified in each panel were from at least 3 independent experiments, which is the same throughout the later figures.

As suggested earlier, the enrichment of the glycolytic enzymes in the propagating waves may concentrate the enzymes and thus enhance the overall glycolytic activity. As a reflection of the glycolytic activity, we utilized the biosensor iATP (mRuby-iATPSnFR) ^33^ to measure the intracellular level of ATP. The ratio of cpGFP to mRuby is correlated with the relative level of ATP (**Figure S2A**). First, we examined the response of this biosensor to inhibitors of glycolysis, 2-Deoxy-D-glucose (2-DDG) and 3-Bromopyruvate (3-BPA), and oxidative phosphorylation (OXPHOS), OAR respectively (**Figure S2B**). When DB was added, there was a major drop of signal in cpGFP but little change in mRuby (**Figure S2C** **and Video S10**). The ratio of cpGFP to mRuby decreased ∼70% in less than 10 min after addition of DB (**Figure S2D**). Later application of OAR did not further reduce ATP (**Figure S2D,** see also **Video S10**). When OAR was applied first, the ratio of cpGFP to mRuby was reduced by less than 10%, while further application of DB reduced the ratio by ∼60% (**Figure S2E**). These experiments suggest that in MCF-10A M3 cells, ATP is largely produced from glycolysis rather than from OXPHOS, which is consistent with previously published results ^34^. Mitotracker showed that mitochondria were not localized in these waves and protrusions (**Figure S3**).

We used this ATP biosensor to measure the glycolytic activity in response to the EGF and insulin. To focus on the ATP change from glycolysis, we pretreated the cells in OAR to eliminate the ATP contribution from OXPHOS. When EGF and insulin were added, there was a burst of ATP production (**Figure 3G**). The cpGFP/mRuby ratio showed a ∼20% increase within 30 min (**Figures 3H** **and S4A)**. This timing was consistent with the increase in aldolase and PFK associated wave activities as indicated in **Figure 3A-F**. This finding suggests that the induced recruitment of the glycolytic enzymes concentrates them into the waves which increases glycolytic activity and leads to increased ATP production.

We further examined the correlation of the ATP level and the wave activity by blocking the cortical association of the glycolytic enzymes. Latrunculin A (Lat A) eliminated the cytoskeletal waves and resulted in a 20% drop in the basal cpGFP/mRuby ratio (**Figure 3I-K** **and** **Figure S4B**). Subsequent addition of EGF and insulin did not increase the cpGFP/mRuby ratio (**Figures 3I, 3L**, **and** **S4C**). Under this condition, aldolase and PFK did not appear to redistribute to the membrane (**Figure 3M**). Lat A did not completely block the activation of PI3K by EGF and insulin as indicated by membrane recruitment of PH-AKT (**Figure 3M**), suggesting that PI3K activation is not sufficient for enhancing glycolysis (**Figure 3I** and **3L**). However, upon the inhibition of PI3K, the cpGFP/mRuby ratio dropped by ∼25% (**Figure 3N** **and** **3P**). The decrease in glycolytic activity by PI3K inhibition was confirmed with pyruvate and NADH/NAD+ sensors ^35,36^ (**Figures 3O, and S4D-H, Videos S11-12**). These results are quantified in **Figures 3P** and **3Q.** Notably, there was also a ∼50% drop in the wave activity (**Figure 3R**). These results suggest glycolysis may be enhanced by the waves of coupled signaling and cytoskeletal activities.

### Relationship between glycolytic wave activities and cell motility

To investigate the relationship between waves of glycolytic enzymes and cell behavior, we altered the wave activities with synthetic biology and drug treatment. It was previously shown that abruptly lowering plasma membrane PI(4,5)P2 by recruitment Inp54p using chemically induced dimerization (CID) (**Figure S5A**) initiates coordinated PIP3 and F-actin waves and accompanied protrusive activities in relatively quiescent MCF-10A cells ^26^. As shown in **Figure 4A,** upon PI(4,5)P2 reduction, increased aldolase was associated with the increased propagating waves and protrusions, which spiraled around the cell perimeter (**Figures 4B** and **4C**; see also **Figure S5B** and **Video S13**). This observation is similar to the EGF and insulin induced increase in cortical waves, protrusions, and associated aldolase and PFK. Thus, the propensity of the glycolytic enzymes to associate with the cortical waves, and increase ATP production, is strongly correlated with the overall excitability of the cell and the generation of F-actin-based protrusions.

**Figure 4.**
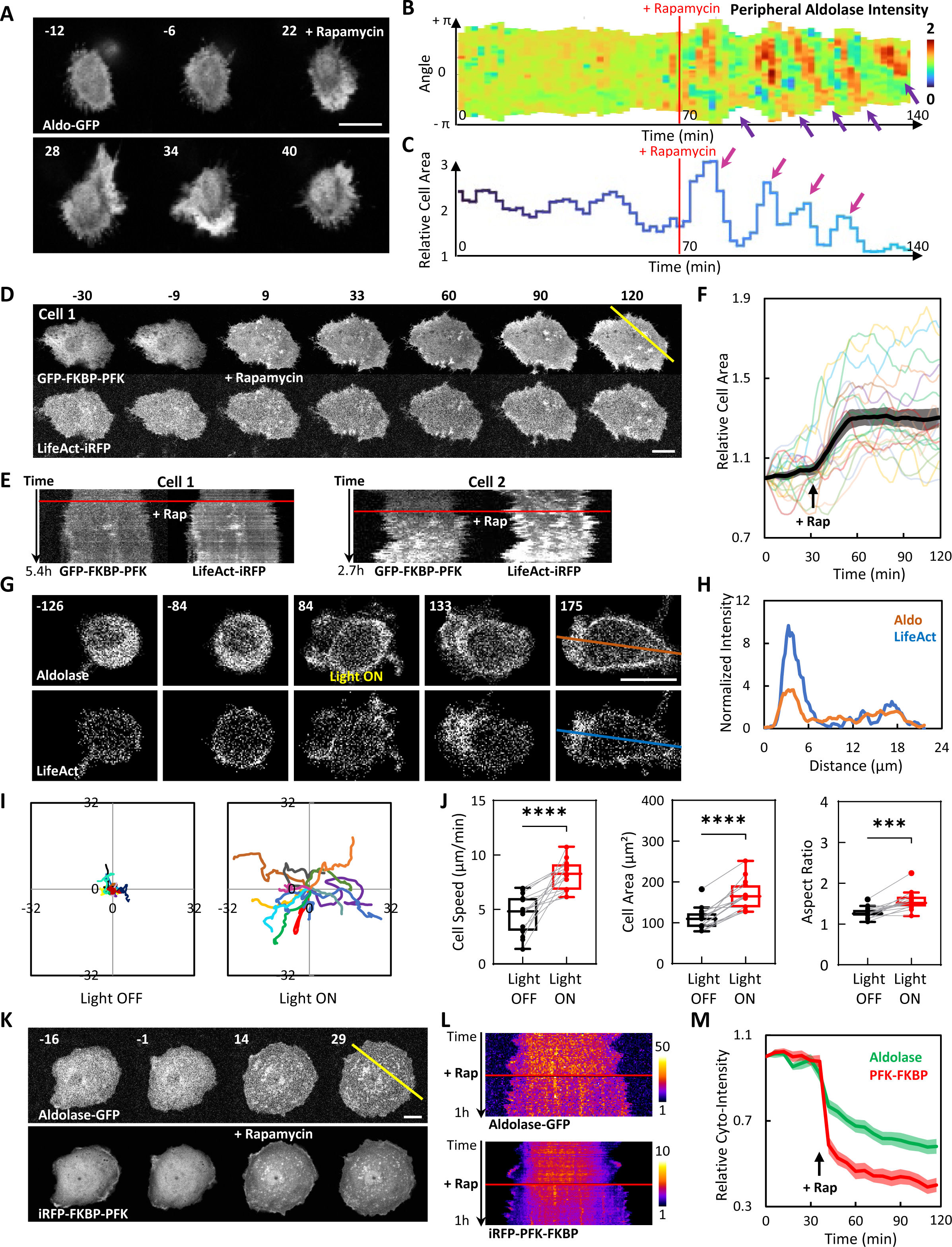
Effects of synthetic and optogenetic perturbations of glycolytic waves on cell motility. (A) Confocal images of the Aldolase-GFP in an MCF-10A M1 cell expressing aldolase-GFP, CFP-Lyn-FRB, and mCherry-FKBP-Inp54p treated with 1 μM rapamycin at 0 min (also see **Video S13**). Schematic explanation of the design of the chemically inducible dimerization (CID) system is shown in **Figure S5A**. More examples of cells are shown in **Figure S5B**. (B) Kymograph of aldolase-GFP signal around the perimeter of the cell in (A) over time. The purple arrows indicate enrichment of aldolase in the rhythmically spiral waves around the cell perimeter. (C) Quantification of the area change of the cell in (A). The pink arrows indicate that the cell area change aligns with enrichment of aldolase in the rhythmically spiral waves around the cell perimeter. (D) Confocal images of an MCF-10A M3 cell expressing GFP-FKBP-PFK, Lyn-FRB, and LifeAct-iRFP treated with 1 μM rapamycin at 0 min (also see **Video S14**). (E) Kymographs of the yellow line in (D) scanning through the cell expressing GFP-FKBP-PFK, Lyn-FRB, and LifeAct-iRFP before and after treatment with 1 μM rapamycin over 5.4 h. An additional example from another cell is shown on the right panel. (F) Quantification of normalized cell area (mean ± SEM) of n = 20 cells in (D) over time. Tracks of changes in individual cells are shown in the color lines. (G) Time lapses confocal images of a differentiated HL-60 neutrophil expressing CRY2PHR-mCherry-Aldolase and LifeAct-miRFP703, before and after 488 nm light illumination (Also see **Video S15**). Time in sec; scale bar: 10 μm. 488 nm Light is turned on at 0 second. Schematic explanation of this experiment is shown in **Figure S6A**. (H) Intensity of aldolase and LifeAct across the orange and blue lines in (G). (I) Centroid tracks of differentiated HL-60 cells showing random motility before and after global recruitment of aldolase. Each track lasts 3 minutes and was reset to the same origin. n = 15 from at least 3 independent experiments. (J) Box-and-whisker plots of HL-60 average cell speed, cell area, and aspect ratio, before (black) and after (red) aldolase recruitment. n = 15 from at least 3 independent experiments. ∗∗∗p ≤ 0.001; ∗∗∗∗p ≤ 0.0001 (Wilcoxon-Mann-Whitney rank-sum test). Quantifications of non-recruitment control are shown in **Figures S6 B-C**. (K) Time lapses confocal images of a cell expressing aldolase-GFP, Lyn-FRB, and iRFP-FKBP-PFK treated with 1 μM rapamycin at 0 min. An additional example from another new cell is shown in **Figures S8A-B**. Also see **Video S16**. (L) Kymographs of the cell in (K) across the yellow line. (M) Plot of normalized intensity of cytosolic aldolase-GFP (mean ± SEM) and cytosolic iRFP-FKBP-PFK (mean ± SEM) of n = 19 cells expressing aldolase-GFP, Lyn-FRB, and iRFP-FKBP-PFK before and after treatment of rapamycin over time. Data of these 19 individual cells are plotted in the color heat map in **Figures S8C-D**. Scale bars are 20 μm for all except (A) and (G) (10 μm). Cells in all panels are MCF-10A M3 cells except (A) (MCF-10A M1 cell) and (G) (HL-60 cell).

We next developed CID and optogenetic systems to rapidly recruit the glycolytic enzymes to the membrane and assess changes in cell behaviors. Surprisingly, in MCF-10A M3 cells, recruitment of PFK to the plasma membrane by CID triggered *de novo* F-actin wave formation and cell spreading (**Figure 4D-4F** **and Video S14**). To demonstrate that this effect was not limited to one enzyme in one cell type, we created a light-inducible recruitment system for aldolase in neutrophil-like HL-60 cells (**Figure S6A**). Quiescent HL-60 cells became polarized and highly motile after the recruitment of aldolase to the membrane (**Figure 4G** **and Video S15**). The recruited aldolase was initially uniformly localized and became enriched in the LifeAct labeled protrusions as the cell became polarized and started to migrate persistently (**Figures 4G, 4H** **and Video S15**). As shown in **Figures 4I** **and** **4J**, migration speed, cell spreading, and polarity increased with aldolase recruitment. These changes were not detected in the non-recruitment controls (**Figure S6B-C**). Together, these results showed the surprising observation that relocalization of a single glycolytic enzyme from the cytosol to the plasma membrane can dramatically alter cell morphology and increase cell migration.

We next explored the effect of inhibition of metabolic pathways on cell migration. As demonstrated above, glycolysis inhibitors abolished the majority of ATP production in the cell while OXPHOS had a minor effect (**Figure S2**). Accordingly, we analyzed the effects of inhibitors of glycolysis and OXPHOS on cell migration. Similar to the drop in cpGFP/mRuby ratio of iATP, the actin wave activities (**Figures S7A, B**), dynamic morphological changes (**Figures S7C-G**) and cell migration tracks (**Figure S7H**) all dramatically decreased upon the inhibition of glycolysis, while further addition of OXPHOS inhibitor did not cause further change. Conversely, addition of OXPHOS inhibitors first caused little change in wave activity and further application of glycolysis inhibitors drastically reduced it (**Figure S7I**), which parallels the ATP change (**Figure S2E**).

How can recruitment of a single enzyme to the plasma membrane enhance glycolysis? To further investigate this surprising result, we examined how membrane recruitment of PFK affects aldolase localization by co-expressing aldolase-mCherry in cells carrying the PFK recruitment system. Consistent with the above findings, recruitment of PFK by CID caused cell spreading and an enhanced level of dynamic protrusions at the perimeter, indicative of increased cortical wave generation (**Figure 4K**). Remarkably, aldolase was also recruited to the plasma membrane (**Figures 4K-M** **and S8; Video S16**). The coordinated behavior of the two enzymes could indicate that PFK recruitment generated cortical waves and aldolase was recruited to them or, alternatively, the enzymes are in a complex. In either case, the coordinated recruitment of glycolytic enzymes can explain why the recruitment of one glycolytic enzyme is sufficient to enhance waves and protrusive activities and ATP production from glycolysis.

### Enhanced glycolytic waves explain energy shift in cancer cells

The cartoon in **Figure 5A** summarizes our conclusions so far indicating that the association of glycolytic enzymes with traveling waves enhances local glycolysis activities, which in turn leads to the formation of further waves. We showed previously that oncogenic transformation with Ras leads to an increased number of dynamically active cortical waves ^26,31^. Cancer cells depend more on glycolysis as an energy source, a phenomenon known as the Warburg effect. Taken together, our studies show that the enrichment of the glycolytic enzymes in the cortical waves provides a large fraction of the ATP in cells. Thus, an increased number of waves, and associated glycolytic enzymes, in transformed cells might underlie the Warburg effect.

**Figure 5.**
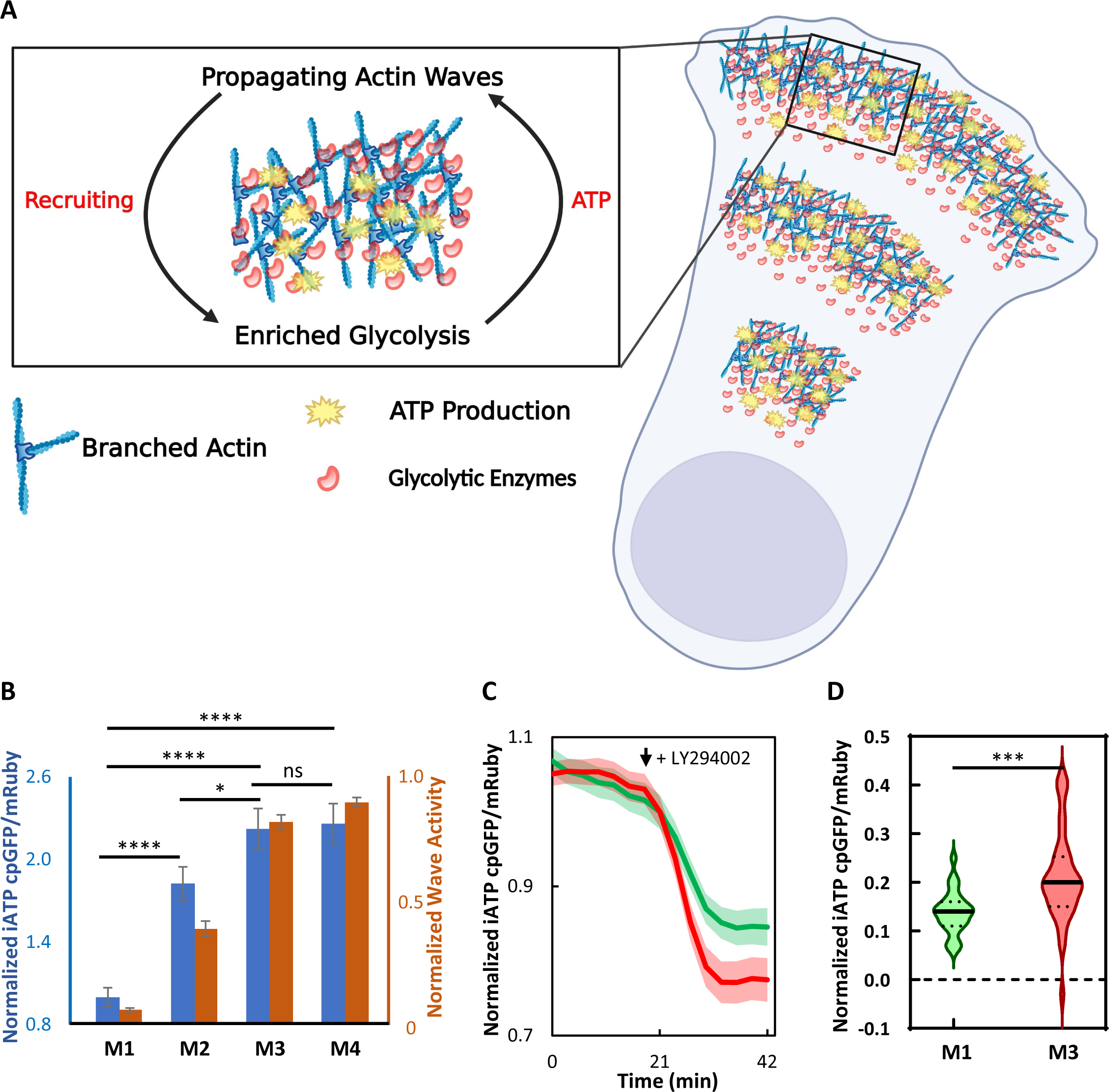
Model and implications in cancer. (A) Model: Enrichment of glycolytic enzymes in self-organized F-actin waves enhances glycolysis to provide energy for migration and other cellular processes. (B) Quantification of cpGFP/mRuby ratio (blue bars) in M1-M4 MCF-10A cells (mean ± SEM of 112 M1 cells, 139 M2 cells, 200 M3 cells, and 198 M4 cells from 5 independent experiments). Unpaired t test with Welch’s correction, ****p < 0.0001, *p < 0.05, ns no significance. Quantification of wave activity from previous published work ^26^ is shown for comparison (orange bars, fraction of cells with waves during a 2-h imaging window, mean ± SEM of 501 M1 cells, 608 M2 cells, 234 M3 cells, and 302 M4 cells). (C) Normalized iATP cpGFP/mRuby ratio signal shown in (mean ± SEM) of n= 27 M1 cells (green) and n= 38 M3 cells (red) treated with LY294002 at the indicated time. The cpGFP/mRuby values are normalized to the time point that LY 294002 was added. Cells were pretreated in OAR before the treatment of LY294002. (D) Average decreases in iATP cpGFP/mRuby ratio before and after LY 294002 treatment in these 27 M1 cells (green) and 38 M3 cells (red) in (C) were shown as violin plots and compared by unpaired t test with Welch’s correction, ***p = 0.0001.

To test the idea, we took advantage of a series of MCF10A derived cell lines, M1-M4, marked by increased oncogenic and metastatic potential. We had previously shown the increasing metastatic potential in this series correlates closely with an increase in cortical wave activities ^26^. We expressed the iATP biosensor in M1-M4 cells. As shown in **Figure 5B**, the M1-M4 series of cells with increasing metastatic potential showed an increasing level of ATP as reflected by cpGFP/mRuby ratio. Given that inhibition of PI3K significantly abrogates wave activity (**Fig 3R**), we next treated M1-M4 cells with the PI3K inhibitor. Inhibiting the cortical waves led to a ∼25% decrease in the cpGFP/mRuby ratio of iATP signal in M3 cells but only ∼15% decrease in M1 cells (**Figure 5C**). That is, inhibition of the cortical waves causes a more significant decrease in ATP production in the more metastatic cells relative to the parentals (**Figure 5D**). Thus, the progressively augmented cortical wave activity, and implied greater cortically associated glycolytic enzymes, seen in the M1-M4 series may provide a mechanism for the greater reliance of the more metastatic cells on ATP from glycolysis.

## Discussion

Our study was prompted by our unexpected discovery that aldolase is associated with propagating cortical waves of Ras/PI3K and F-actin on the cell membrane. Following this up, we found that all of the glycolytic enzymes we had tested are associated with the cell cortex, co-localized and traveling with the signal transduction and cytoskeletal components. Growth factors and other stimuli coordinately increase wave activity and recruitment of glycolytic enzymes into the waves while inhibitors that abolish waves send the enzymes back to the cytosol. Consistently, ATP levels are strongly correlated with the augmentation or abrogation of the waves under these various perturbations. Recruitment of a single glycolytic enzyme induces epithelial cell spreading and, in neutrophils, enhances polarity and migration while inhibition silences membrane undulations. Furthermore, recruitment of PFK causes a co-recruitment of aldolase to the membrane. Importantly, we found an increased frequency of waves and a concomitant increased level of ATP in more metastatic cells, indicating that the higher abundance of waves of glycolytic activity observed in cancer cells may explain the elevated metabolic activity associated with the Warburg effect.

Although glycolysis is traditionally thought to occur in the cytosol, in addition to our report, other studies have suggested alternate localizations or configurations of some glycolytic enzymes. These studies do not necessarily conflict, and may be consistent, with our findings reported here. For example, the aldolase released from stress fibers by PI3K activation may free it up to associate with cortical waves, although we saw no evidence for this. However, our findings add an unexpected dimension to the understanding of glycolysis, as this work clearly suggests a link to cell migration, since the glycolytic enzymes associate with the very waves that underlie protrusion formation.

Our observations raise several questions about the molecular basis and consequences of glycolytic waves. How are the enzymes recruited to the waves? Several glycolytic enzymes have been shown to bind F-actin *in vitro* ^6,7,37–40^. The molecular determinant of binding has not been clearly elucidated for the majority of these enzymes, although an actin binding region was identified in aldolase ^8^. However, the width of the glycolytic bands is wider than those of the LifeAct, suggesting that the localization of glycolytic enzymes may not be through direct binding to newly polymerized actin or actin at all. Instead, these enzymes may be associated with other molecules such as signaling molecules, which display similar diffusive wave bands. Importantly, synthetic recruitment of PFK caused a co-recruitment of aldolase. This may suggest a direct association of the enzymes with each other or association with a common scaffold.

What is the effect of the enzymes associating with the waves on the rate of glycolysis? High concentrations of glycolytic enzymes in these waves may enhance the reaction rate. Moreover, proximity of the substrate, produced from the previous glycolytic reaction, to the glycolytic enzyme could also lead to more efficient glycolysis. Based on our imaging studies at least 15% of the enzymes are recruited into the waves. We did not have the resolution to determine whether the enzymes were associated with the membrane or with the cortex. If a cytosolic protein were concentrated into a 0.5 μm shell covering the entire cortex of a 30 μm cell, the concentration increase would be 20-fold. Although only 15% of the enzymes are recruited, they are recruited into a small area of the cortex covered by waves, so that the concentration effect is quite significant. If the enzymes actually associate with the membrane rather than the cortex, the concentration effect could be much higher.

What is the functional significance of the glycolytic waves? Our study suggests that local production of ATP may provide the energy that fuels cortical wave formation. Existing studies have shown that *Dictyostelium*, oocytes, mast cells, epithelial cells, neurons, and other cells display these cortical waves, which regulate many biological processes such as cell growth, cell cycle, phagocytosis, protein trafficking, synaptogenesis, and migration in these cells ^41–50^. The glycolytic waves are likely to control these functions. While we clearly showed that the association of the enzymes with the cortical waves is correlated with ATP levels and cellular protrusions/motility, our imaging studies did not reveal a localized production of ATP. Further studies are needed to resolve this issue.

Our study provides an explanation for the Warburg effect. We have previously suggested that cancer cells are shifted to a lower threshold “state” of key signal transduction and cytoskeletal networks, such as the Ras/PI3K/ERK network involved in cellular transformation. Our current study shows that oncogenic transformation leads to increased glycolytic waves, leading to higher glycolytic activity. Thus, the increased association of the glycolytic enzymes with the cortex accompanies a shift in the “state” of the excitable networks. Tumor cells often reprogram their energy metabolism by using glycolysis as the main source of ATP production rather than oxidative phosphorylation, even when oxygen is available ^51–53^. This effect, known as aerobic glycolysis or Warburg effect, was reported by Otto Warburg nearly 100 years ago ^3,4^, yet the mechanism is not fully understood. The propensity of the glycolytic enzymes to associate with the cortical waves would provide an elegant mechanism for coordinating glycolytic activity with the state of the excitable networks in normal and oncogenic transformed cells.

## Materials and Methods

### Cells

M1 (MCF-10A), M2 (MCF-10AT1k.cl2), M3 (MCF-10CA1h), and M4 (MCF-10CA1a.cl1) cells, purchased from the Animal Model and Therapeutic Evaluation Core (AMTEC) of Karmanos Cancer Institute of Wayne State University, were all grown at 37°C in 5% CO2 using DMEM/F-12 medium (Gibco, #10565042) supplemented with 5% horse serum (Gibco, #26050088), 20 ng/ml EGF (Sigma, #E9644), 100 ng/ml cholera toxin (Sigma, #C-8052), 0.5 mg/ml hydrocortisone (Sigma, #H-0888) and 10 μg/ml insulin (Sigma #I-1882). Human neutrophil-like HL-60 cells were gifted by Orion Weiner (UCSF) and grown in supplemented RPMI medium 1640 (Gibco #22400-089) as described previously ^28,54^. For cell differentiation, neutrophils were incubated with 1.3% DMSO for 5-7 days before experimentation ^28,54^.

Transient transfections of the cells were performed using Lipofectamine 3000 (Invitrogen, #L3000008) following manufacturer’s instructions. Cells were transferred to 35 mm glass-bottom dishes (Mattek, #P35G-0.170-14-C) or chambered coverglass (Lab-Tek, #155409PK) and allowed to attach overnight prior to imaging. Cells were seeded and incubated at 37°C in 5% CO2 overnight before live cell imaging. For EGF and Insulin stimulation assays, MCF-10A (M1 - M4) cells were starved in pure DMEM/F-12 medium for 24 hours before stimulation.

### Plasmids

Constructs of CFP-Lyn-FRB, and mCherry-FKBP-Inp54p were obtained from Inoue Lab (JHU). GFP/RFP-PH-AKT and RFP-LifeAct were obtained from Desiderio Lab (JHU). Aldolase-GFP was generously provided by the Wulf Lab (Harvard) 10. PFK-GFP (#116940) 11, Truncated-HK-GFP (#21918) 55, Lifeact-iRFP (#103032) 56, mRuby3-iATPSnFR1.0 (#102551) 33, PyronicSF-mRuby (#124830) 36, and Peredox-mCherry (#32380) 35 constructs were obtained from AddGene. Enolase-RFP, PK-RFP, GAPDH-RFP, and GFP/iRFP-FKBP-PFK were generated in this study.

Lyn-FRB and FKBP-Inp54p were subcloned into the lenti-viral expression plasmid pFUW2. 3rd generation lentiviral constructs, CIBN-CAAX/pLJM1 and LifeAct-miRFP703/pLJM1, were generated in a previous study28. The human AldolaseA ORF (1092 bases) was PCR-amplified and cloned into BspEI/SalI sites of the PiggyBac™ transposon plasmid to generate the CRY2PHR-mCherry-AldolaseA/pPB construct. All constructs were verified by sequencing at the JHMI Synthesis and Sequencing Facility.

### Drugs

Stocks of 25 mM Latrunculin A (Enzo, #BML-T119-0100), 50 mM LY294002 (Invitrogen, #PHZ1144), 10mM Oligomycin (Cell Signaling Technology #9996L), 5 mM Antimycin A (Sigma, #A8674), 5 uM Rotenone (Sigma, #R8875), 400mM 2-Deoxy-D-Glucose (BioVision, #B1048), 400mM 3-Bromopyruvic acid (BioVision, #B1045), 20 mM MitoTracker (Thermo Fisher, #M7514) and 10 mM Rapamycin (Cayman, #13346) were prepared by dissolving the chemicals in DMSO. The stocks were diluted to the indicated final concentrations in culture medium or live cell imaging medium. The EGF stock solution was prepared by dissolving EGF (Sigma, #E9644) in 10 mM acetic acid to a final concentration of 1 mg/ml. Insulin (Sigma #I-1882) was resuspended at 10 mg/ml in sterile ddH2O containing 1% glacial acetic acid. Hydrocortisone (Sigma #H-0888) was resuspended at 1 mg/ml in 200 proof ethanol. Cholera toxin (Sigma #C-8052) was resuspended at 1 mg/ml in sterile ddH2O and stored at 4°C. All drug stocks except cholera toxin were stored at −20°C.

#### Virus generation

Cell Seeding and Transfection: On day 1, 293T cells were seeded at a density of 6x10^5 cells/ml in 25 ml of culture medium into 15 cm cell culture dishes. The next day, conventional calcium phosphate transfection was employed to introduce expression and packaging plasmids into the 293T cells. The transfection mixture consisted of 20 µg of pFUW2, 9.375 µg each of pMDL, pRSV, and pCMV plasmids, combined with 250 µl of CaCl2 and sterile deionized water, bringing the total volume to 2.5 ml. This mixture was then combined with 2.5 ml of 2x HEPES buffer (pH 7.05) and incubated for 5 minutes. The resulting transfection mixture was gently added to the plated cells. After 4-6 hours, the medium was replaced with fresh culture medium.

Virus Collection and Concentration: On day 5, the medium containing viral particles was harvested from the transfected cells. It was initially centrifuged at 1000 rpm for 3 minutes to remove cellular debris and then filtered through a 0.45 µm filter. Subsequently, the filtrate was subjected to ultracentrifugation at 25,000 rpm for 90 minutes at 4°C using a Beckman ultracentrifuge. The supernatant was discarded, and the viral pellet was resuspended in 70 µl of phosphate-buffered saline (PBS) and incubated overnight at 4°C for virus concentration. The concentrated virus was aliquoted into 25 µl portions and stored at −80°C.

### Stable line generation

A stable HL-60 cell line co-expressing CIBN-CAAX and LifeActmiRFP703 was generated using a lentiviral-based approach described in previous studies ^23,28^. In this dual expressing cell line, we stably expressed CRY2PHR-mCherry-AldolaseA via transposon integration ^23,28,54^.

### Microscopy

Confocal microscopy has been described previously ^26^. Briefly, confocal microscopy was carried out on Zeiss AxioObserver inverted microscope with either LSM780-Quasar (34-channel spectral, high-sensitivity gallium arsenide phosphide detectors, GaAsP) or LSM880 confocal module controlled by the Zen software. All live cell imaging was carried out in a temperature/humidity/CO2-regulated chamber.

### Optogenetic experiments

Optical experiments were done without chemoattractant. Photoactivation was performed with a 488 nm excitation Argon laser, CRY2PHR-mCherry-AldolaseA was visualized with a 561 nm excitation solid-state laser, and LifeAct-RFP703 was excited with a diode laser (633 nm excitation). A 40X/1.30 Plan-Neofluar oil objective was used. Pre-treated, differentiated HL-60 cells were prepared for Zeiss LSM780 confocal microscopy on fibronectin-coated chambered coverglass as described earlier ^23,28,54^. For global recruitment, the Argon laser was switched on after imaging for 3 min. Photoactivation and image acquisition was done once every 6-7 seconds. The laser intensity during image capture was maintained at 0.14–0.17 W/cm2 at the objective, which ensured effective AldolaseA recruitment over the cell periphery without inducing photo damage.

### Imaging quantifications

#### Cell Migration and Morphological Analysis

All migration track, cell speed or area, and aspect ratio analyses were carried out by segmenting MCF10A or differentiated HL-60 cells on Fiji/ImageJ 1.52i software ^57^, as described previously ^26, 28^.

### Quantification of Biosensors for ATP (Ruby-iATPSnFR), Pyruvate (mRuby-PyronicSF), and NADH/NAD+ (mCherry-Peredox)

Two channels in each biosensor were captured simultaneously in a confocal microscope. The mask of image was firstly obtained by binarizing the images of mRuby or mCherry channel following despeckling, proper thresholding, and holes-filling in Fiji/ImageJ. The background-removal images were generated by multiplying the images of all channels to the corresponding masks. The Ruby channel of iATP and Pyronic or mCherry of Peredox were added for a very small value to make the denominator non-zero. The ratio images were gained by dividing the background removed cpGFP channel of iATP, Pyronic, or Peredox channel to its processed corresponding Ruby or mCherry channel.

### Statistical Analysis

GraphPad Prism 7 software was used for all statistical analyses. All quantifications are displayed as mean ± SD or SEM. Two-tailed P-values were calculated using a parametric t test. P<0.05 was considered statistically significant. Further details of statistical parameters and methods are reported in the corresponding figure legends.

## Acknowledgements

The authors thank all members of the Devreotes laboratory, Doug Robinson, Pablo Iglesias, Miho Ijima, Halimatu Mohammed and Liang Wang (Janetopoulos Lab), and Hideki Nakamura (Inoue Lab) for helpful discussions and suggestions. We thank Yunlu Li for help in data analysis. We thank Takanari Inoue (JHU), Stephen Desiderio (JHU), and the Gerbug Wulf (Harvard) laboratories for their generous provision of plasmids. We thank Orion Weiner (UCSF) and Sean Collins (UC Davis) for providing HL-60 cells and transposon plasmids, respectively. This work was supported by NIH grant R35 GM118177 (to P.N.D.), AFOSR MURI FA95501610052 (to P.N.D.), DARPA HR0011-16-C-0139 (to P.N.D.), a Cervical Cancer SPORE P50CA098252 Pilot Project Award (to C.H.H.), R01GM136711 (to C.H.H.), the Sol Goldman Pancreatic Cancer Research Center (to C.H.H.), a Johns Hopkins Discovery Award (to C.H.H.), a W.W. Smith Charitable Trust Award (to C.J) and an NIH grant S10 OD016374 (to S. Kuo of the JHU Microscope Facility).

## Author Contributions

H.Z., P.N.D., and C.J. conceived of the overall project. H.Z. designed the experimental plan with assistance from P.N.D., C.H.H., and C.J.. H.Z., D.S.P., and J.B. designed, and J.B generated expression constructs. H.Z. performed all the experiments and data analysis for all MCF10A cells. D.S.P. generated stable cell line, designed and performed imaging experiments, and conducted data analyses in HL-60 Cells (Fig 4G-J, S6). H.Z. prepared the figures and videos with assistance from P.N.D., C.H.H., and C.J.. H.Z., C.J., C.H.H., and P.N.D. wrote the manuscript. P.N.D. supervised all aspects of the project. All authors agreed to the final submitted manuscript.

## Supplemental Figures

**Figure S1.**
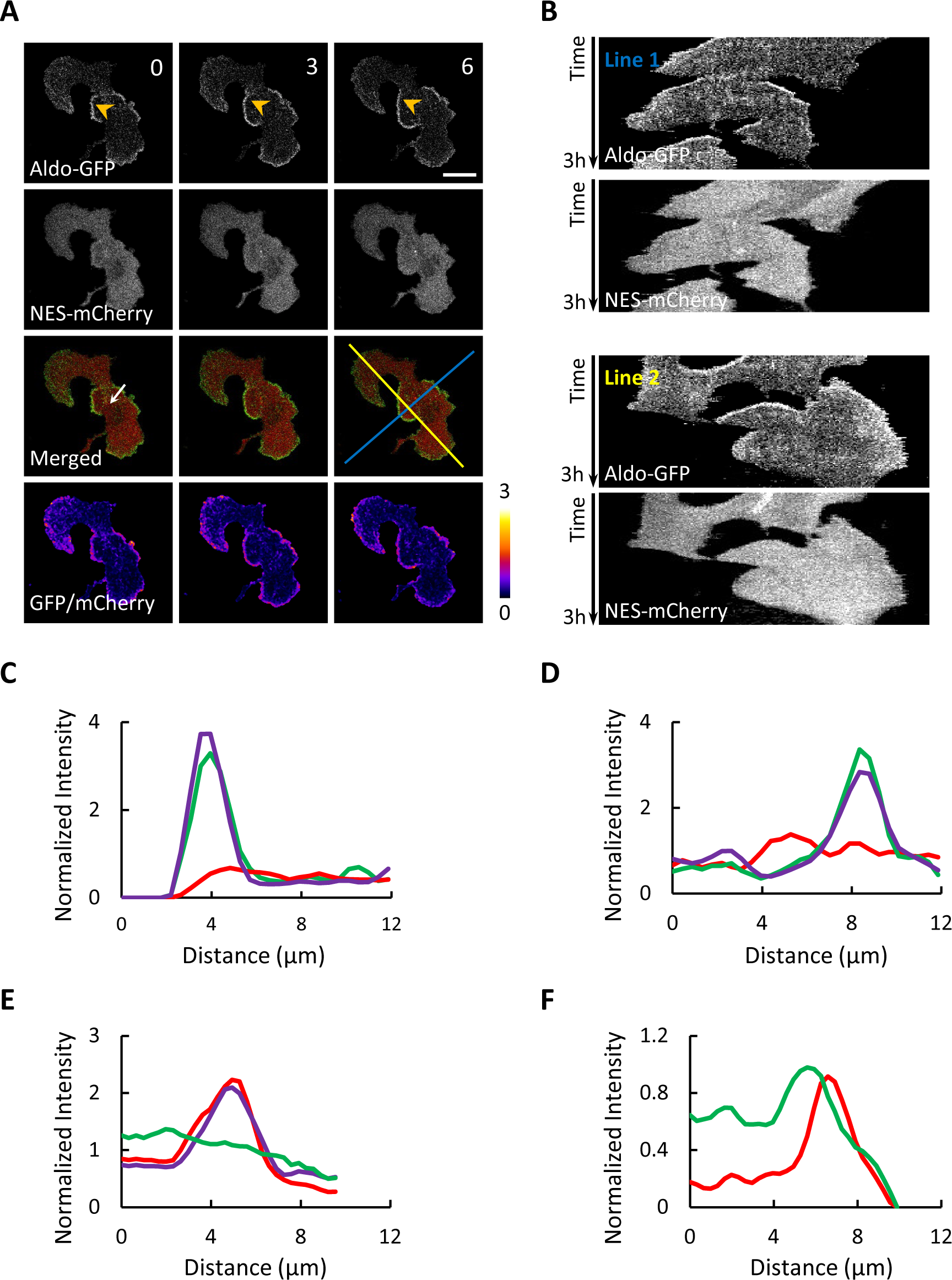
Enrichment of aldolase in F-actin waves and protrusions. (A) Another example of an MCF-10A M3 cell expressing aldolase-GFP and NES-mCherry similar to **Figure 1A**. Scale bar: 20 μm. (B) Kymographs along the two lines in (A) over 3 h. (C) Normalized intensity of aldolase-GFP (green), NES-mCherry (red), and the GFP/mCherry ratio (purple) across the white arrow in (A). (D) Normalized intensity of aldolase-GFP (green), NES-mCherry (red), and the GFP/mCherry ratio (purple) across the white arrow in **Figure 1A**. (E) Normalized intensity of NES-GFP (green), LifeAct-RFP (red), and the RFP/GFP ratio (purple) across the white arrow in **Figure 1C**. (F) Normalized intensity of aldolase-GFP (green) and LifeAct-RFP (red) across the white arrow in **Figure 1E**.

**Figure S2.**
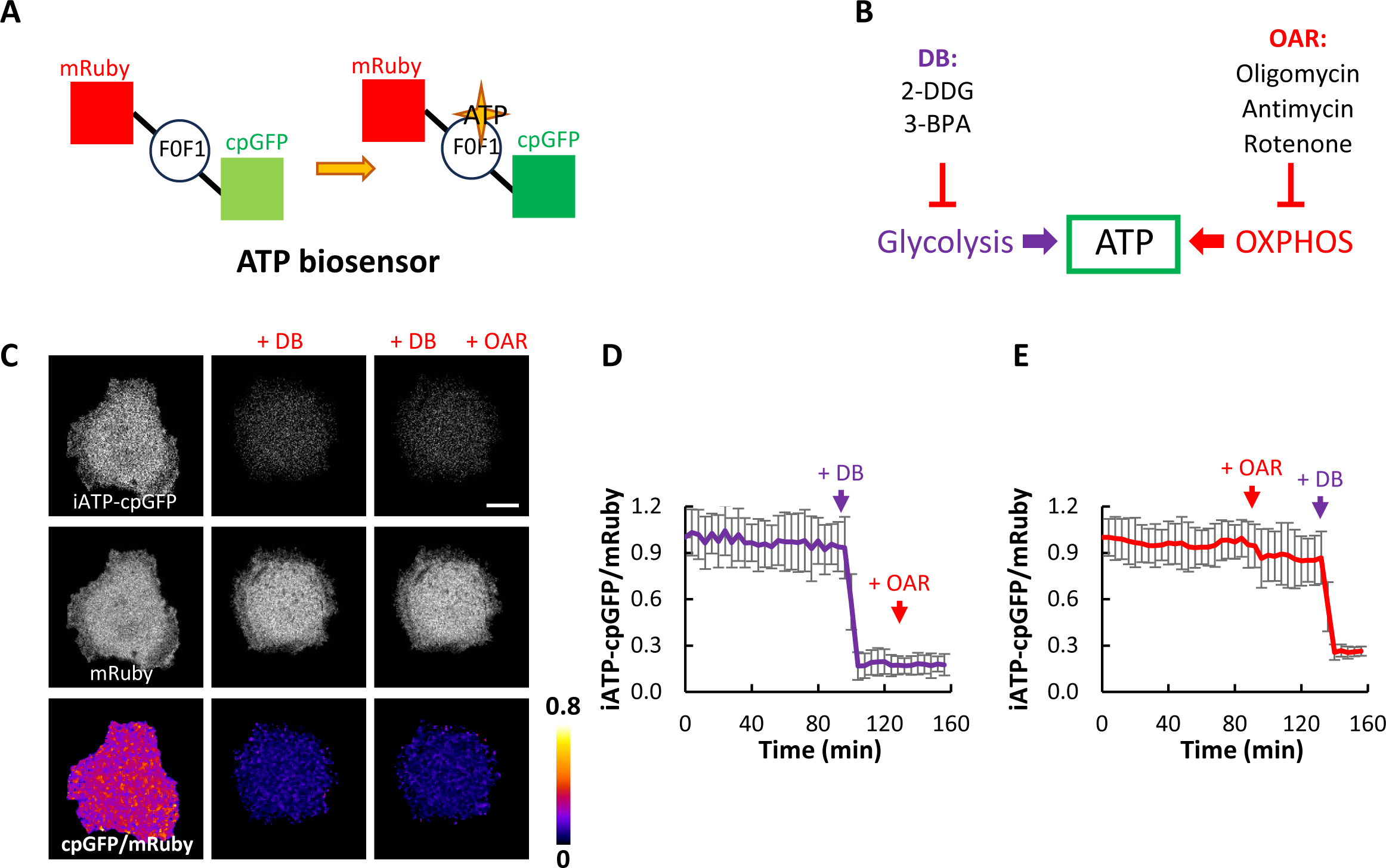
Effects of glycolysis and OXPHOS inhibition on ATP levels. (A) Design of the iATP biosensor ^33^. Binding of ATP to the F0F1-ATPase subunit causes a conformational change of the circularly permuted GFP (cpGFP), leading to increased fluorescence. (B) Inhibitors for glycolysis and OXPHOS used in this study. 2-DDG: 2-Deoxy-D-glucose (10 mM), 3-BPA: 3-Bromopyruvic acid (50 uM), Oligomycin (5 μM), Rotenone (1 μM), Antimycin A (1 μM). (C) Images of cpGFP, mRuby, and the cpGFP/mRuby ratio of a cell expressing the iATP biosensor treated with DB and OAR (also see **Video S10**). Scale bar: 20 μm. (D) Plot of normalized iATP cpGFP/mRuby (mean ± SEM) over time of 15 cells treated with DB followed by OAR. (E) Plot of normalized iATP cpGFP/mRuby (mean ± SEM) over time of 15 cells treated with OAR followed by DB.

**Figure S3.**
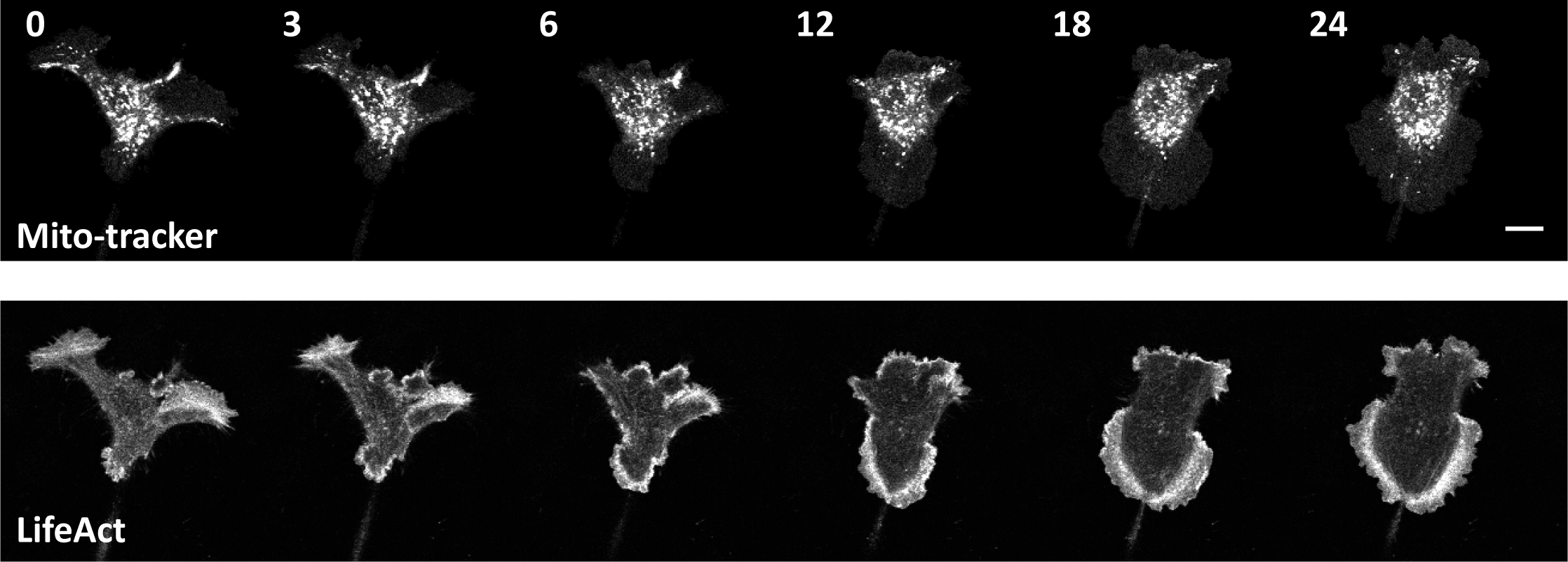
Absence of mitochondria from the waves and protrusions. Confocal images of a cell expressing Mito-tracker-Green and LifeAct-iRFP. Scale bar: 20 μm.

**Figure S4.**
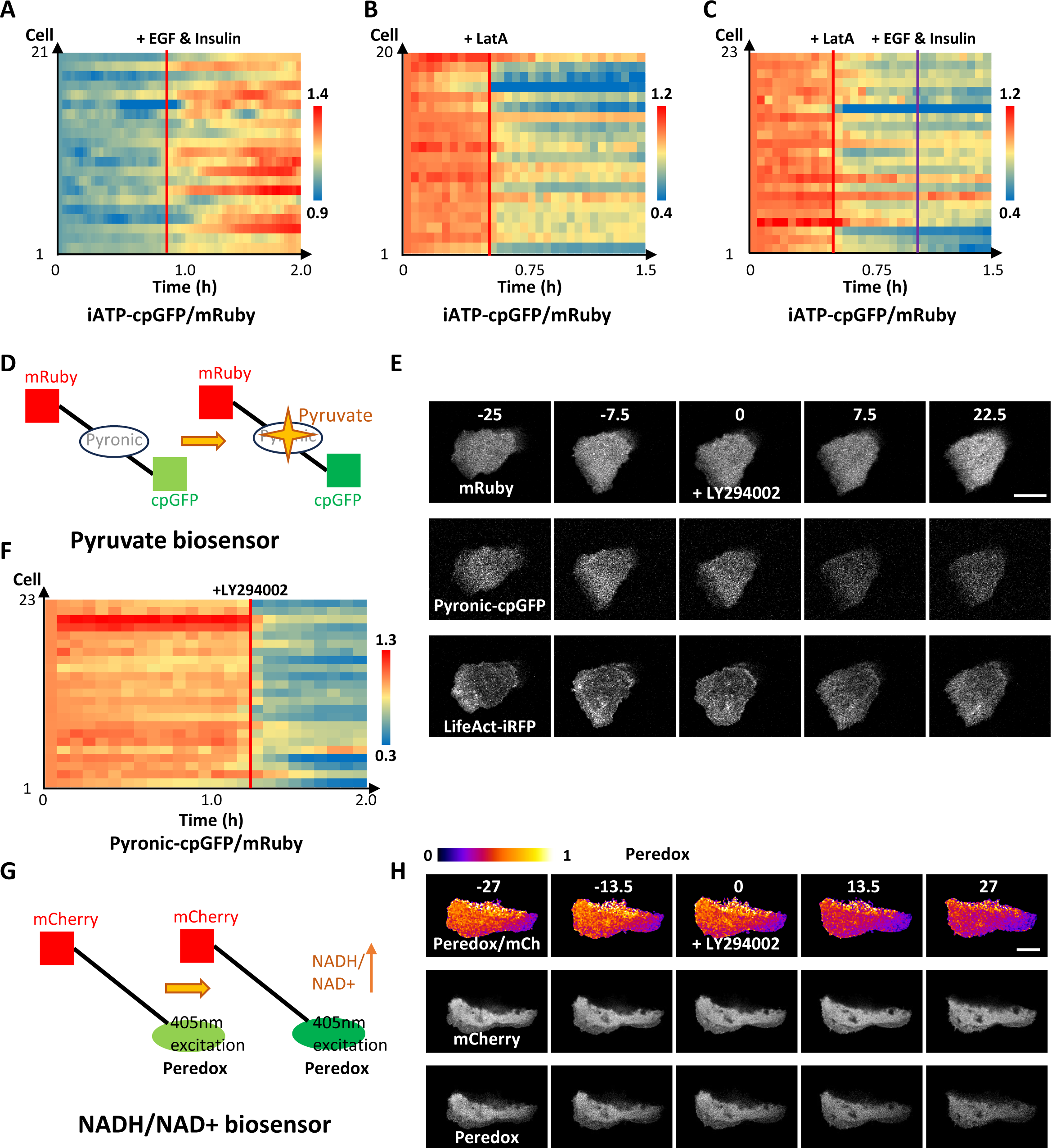
Effects of perturbing wave activities on ATP, pyruvate, or NADH/NAD+ levels. (A) Plots of iATP cpGFP/mRuby ratio in 21 cells stimulated with EGF and insulin in **Figure 3H**. (B) Plots of iATP cpGFP/mRuby ratio in 20 cells treated with Latrunculin A in **Figure 3K**. (C) Plots of iATP cpGFP/mRuby ratio in 23 cells upon treatment with Latrunculin A followed by EGF and insulin in **Figure 3L**. (D) Design of the pyruvate biosensor Pyronic ^36^. Binding of pyruvate causes increased fluorescence of cpGFP. (E-F) (E) Images of a cell expressing Pyronic and LifeAct-iRFP treated with 50 μM LY294002 at the 0 min (corresponding to **Figure 3O,** also see **Video S11**). Scale bar: 20 μm. The responses of 23 cells over time are shown in (F). (G) Design of the NADH/NAD+ biosensor Peredox ^35^. Binding of NADH causes increased fluorescence of circularly permuted T-Sapphire, a GFP variant with peak excitation around 400 nm. (H) Images of Peredox/mCherry ratio, mCherry, and Peredox of a cell expressing the NADH/NAD+ biosensor treated with 50 μM LY294002 at 0 min. Scale bar: 20 μm. Also see **Video S12.**

**Figure S5.**
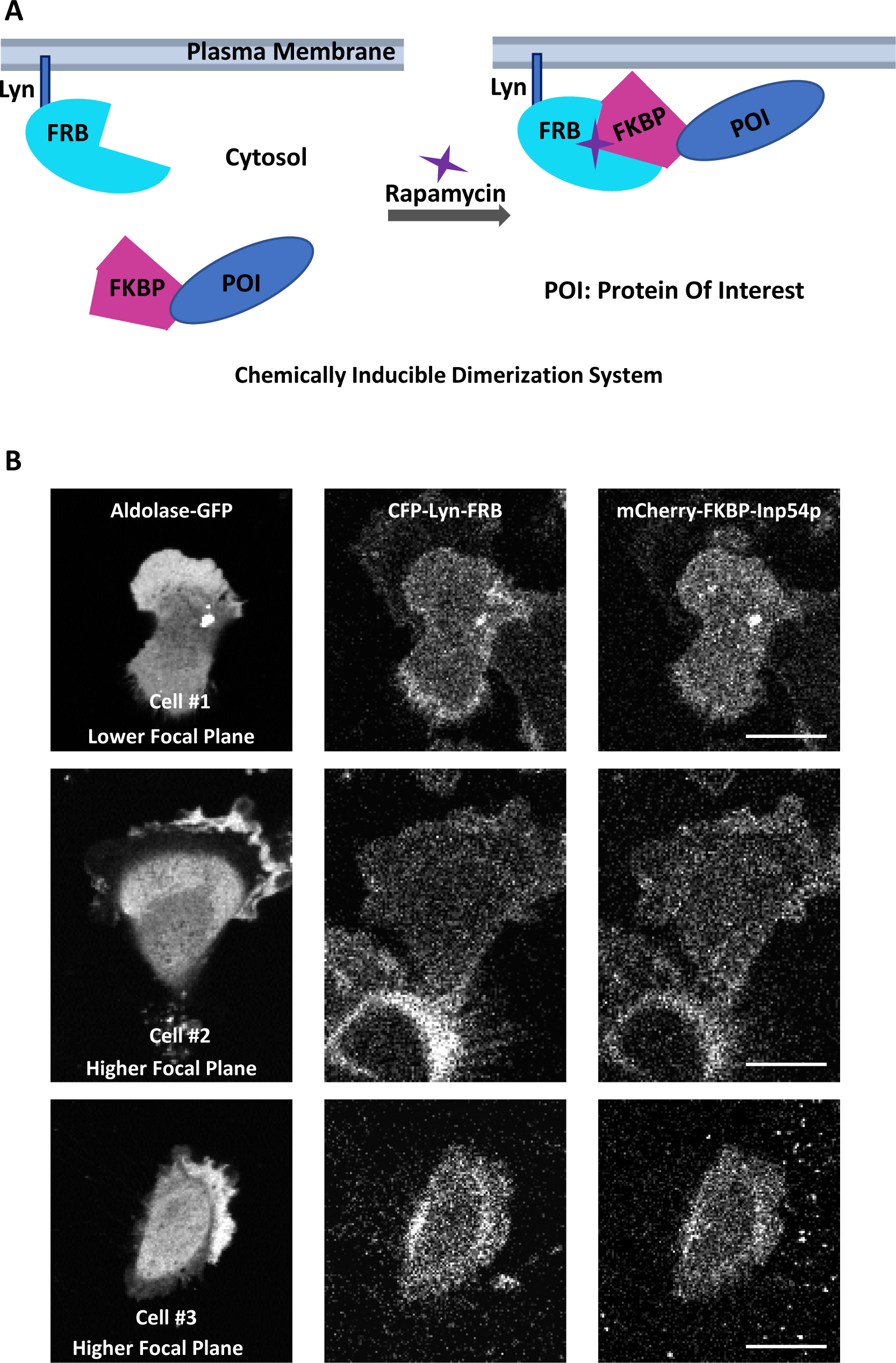
Enrichment of aldolase in waves and protrusions upon PIP2 depletion. (A) Schematic illustration of chemically induced dimerization (CID) used in **Figure 4**. (B) Three more examples of cells showing aldolase-GFP enriched in the spiral peripheral waves and protrusions in MCF10A M1 cells after PIP(4,5)P2 lowering by recruiting the inp54p to the cell membrane. Scale bar: 20 μm.

**Figure S6.**
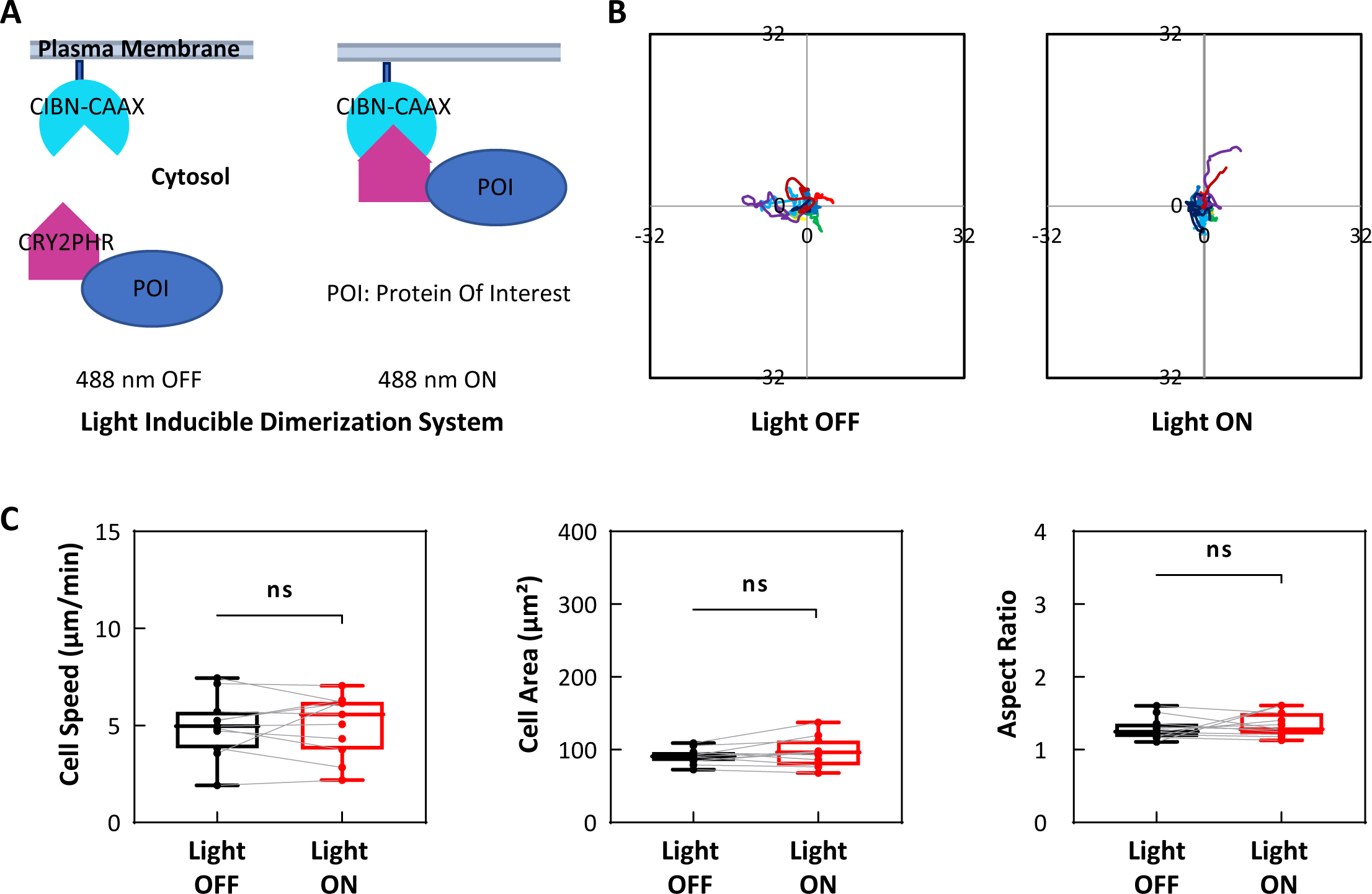
Optogenetic membrane recruitment of aldolase and controls. (A) Schematic illustration of light inducible dimerization system used in **Figures 4G-J**. (B-C) Non-recruitment control of aldolase to the plasma membrane has no effect on migration tracks (B) and cell speed, cell area, and aspect ratio (C) of HL-60 cells, in contrast to the cell in **Figures 4G-J.**

**Figure S7.**
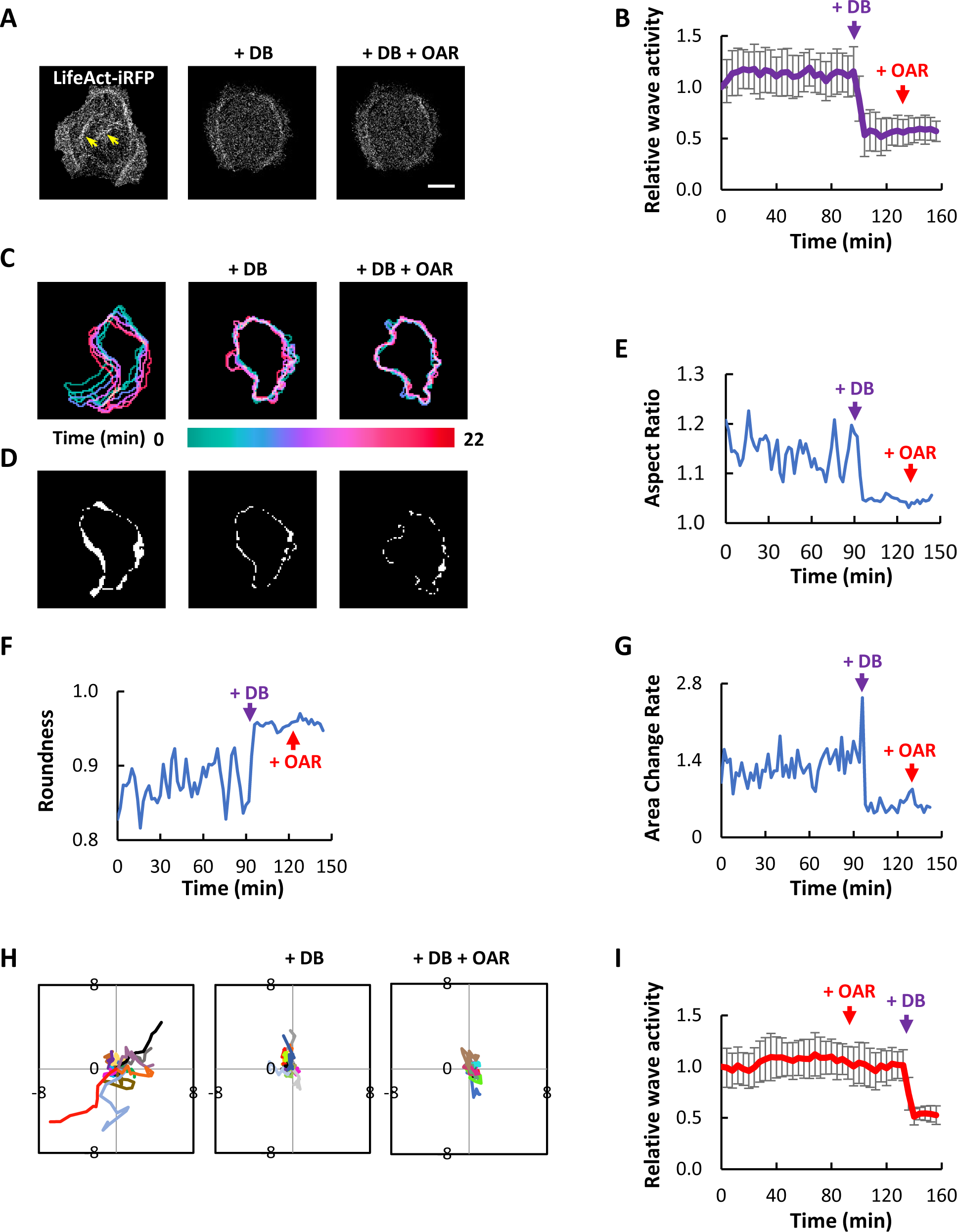
Effects of glycolysis and OXPHOS inhibition on waves and cell morphodynamics. (A) Images of LifeAct-iRFP in an MCF-10A M3 cell treated with DB followed by OAR. (B) Plot of normalized ratio of membrane to cytosol LifeAct (mean ± SEM) over time of 15 cells treated with DB followed by OAR at the indicated time. (C) Color-coded overlays showing the progression of the shape change of a cell over 22 min before and after DB and OAR treatment. (D) Changes in cell morphology between two consecutive frames from the time-lapse images in (C). (E-G) Plot of the aspect ratio (E), roundness (F), and the rate of area change (G) over time for the cell in (A). (H) Centroid tracks of 15 cells showing random motility before and after treatment with DB and OAR. (I) Plot of normalized ratio of membrane to cytosol LifeAct (mean ± SEM) over time of 15 cells treated with OAR followed by DB at the indicated time.

**Figure S8.**
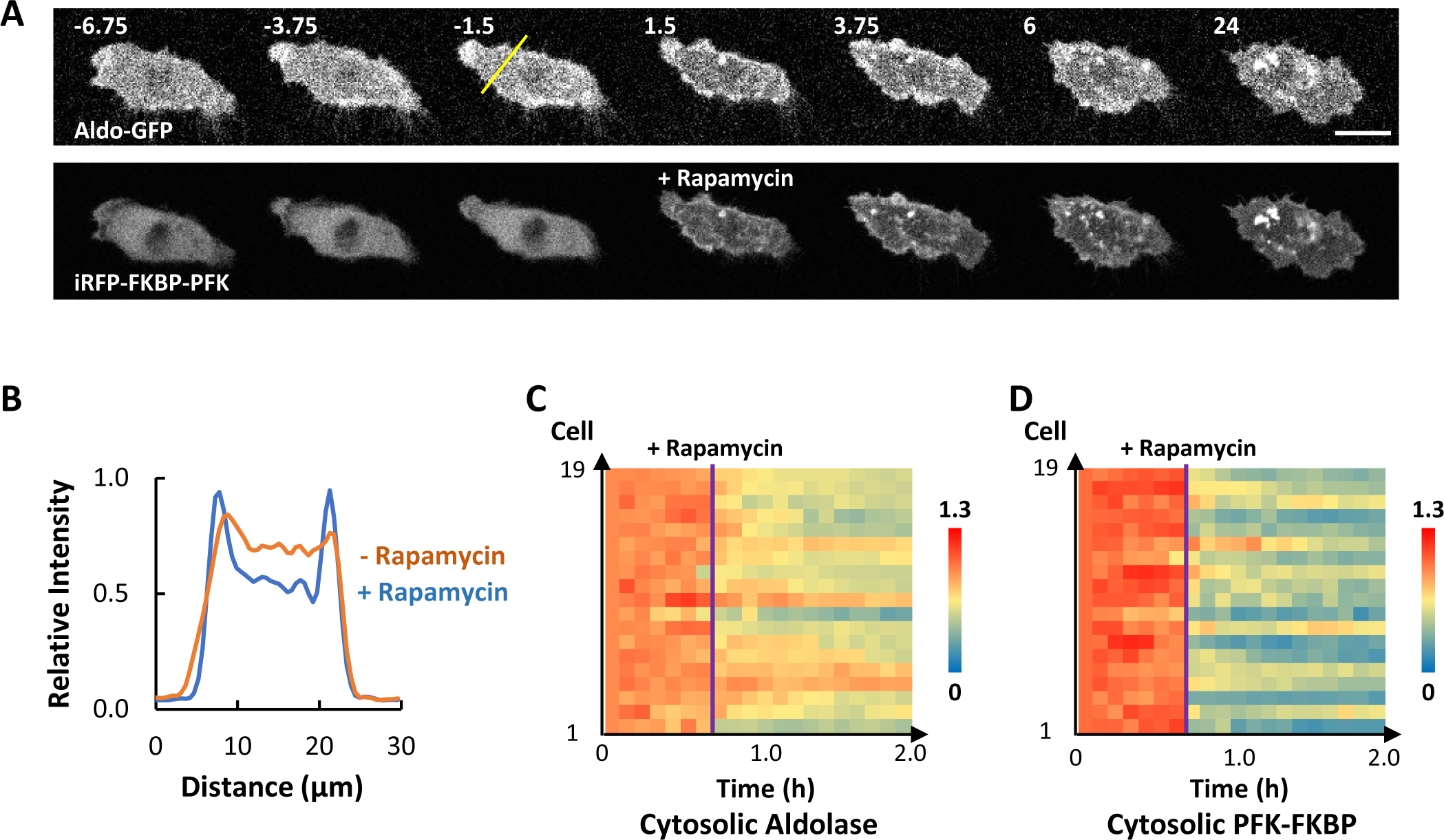
Effect of PFK membrane recruitment on aldolase localization. (A) Time lapses confocal images of a cell expressing aldolase-GFP, Lyn-FRB, and iRFP-FKBP-PFK treated with 1 μM rapamycin at 0 min, similar to the cell in **Figure 4K**. Scale bar: 20 μm. (B) Relative intensity of aldolase-GFP across the yellow line in (A) before and after rapamycin treatment. (C-D) Responses of cytosolic aldolase (C) and cytosolic PFK-FKBP (D) to rapamycin treatment in 19 cells expressing aldolase-GFP, Lyn-FRB, and iRFP-FKBP-PFK (corresponding to **Figure 4M**).

## Supplemental Videos

**Video S1.** Panels from left to right show the aldolase-GFP, NES-mCherry, the merged, and the GFP/mCherry ratio videos of glycolytic waves propagating on the basal surface of an MCF-10A M3 cell. Color scale in ratio images is 0-16. Time stamp in the video is shown as hour:min:sec, which is consistent throughout all the later videos. Individual scale bar is indicated in each video. Related to **Figure 1A**.

**Video S2.** Panels from left to right show the LifeAct-RFP, NES-GFP, the merged, and the RFP/GFP ratio videos of glycolytic waves propagating on the basal surface of an MCF-10A M3 cell. Color scale in ratio images is 0-7. Related to **Figure 1C**.

**Video S3** Panels from left to right show the aldolase-GFP, LifeAct-RFP, and the merged videos of glycolytic waves propagating on the basal surface of an MCF-10A M3 cell. Related to **Figure 1E**.

**Video S4.** Panels from left to right show the hexokinase-GFP, LifeAct-iRFP, and the merged videos of glycolytic waves propagating on the basal surface of an MCF-10A M3 cell. Related to **Figure 2B**.

**Video S5.** Panels from left to right show the phosphofructokinase-GFP, LifeAct-iRFP, and the merged videos of glycolytic waves propagating on the basal surface of an MCF-10A M3 cell. Related to **Figure 2D**.

**Video S6.** Panels from left to right show the GAPDH-RFP, LifeAct-iRFP, and the merged videos of glycolytic waves propagating on the basal surface of an MCF-10A M3 cell. Related to **Figure 2F**.

**Video S7.** Panels from left to right show the enolase-RFP, LifeAct-iRFP, and the merged videos of glycolytic waves propagating on the basal surface of an MCF-10A M3 cell. Related to **Figure 2H**.

**Video S8.** Panels from left to right show the pyruvate kinase-RFP, LifeAct-iRFP, and the merged videos of glycolytic waves propagating on the basal surface of an MCF-10A M3 cell. Related to **Figure 2J**.

**Video S9.** Two Cells on top are expressing aldolase-GFP while the two in the bottom are expressing PFK-GFP. Glycolytic waves propagating on the basal surface of these MCF-10A M3 cells are imaged before and after treatment with EGF and Insulin. Related to **Figure 3A** **and** **3D**.

**Video S10.** Video of the basal surface of an MCF-10A M3 cell expressing iATP-cpGFP, mRuby, the cpGFP/mRuby ratio, and LifeAct-iRFP before and after treatment with DB followed by OAR. Color scale in ratio image is 0-1. Related to **Figure S2C**.

**Video S11.** Video of the basal surface of an MCF-10A M3 cell expressing pyronic-cpGFP, mRuby, LifeAct-iRFP, and the cpGFP/mRuby ratio prior to and after the addition of the PI3K inhibitor LY294002. Color scale in ratio image is 0-1. Related to **Figure S4E**.

**Video S12.** Video of the basal surface and higher focal plane of an MCF-10A M3 cell expressing peredex/mCherry ratio, peredex, and mCherry prior to and after the addition of the PI3K inhibitor LY294002. Color scale in ratio image is 0-1. Related to **Figure S4H**.

**Video S13.** Video showing the recruitment of aldolase-GFP (channel shown in the video) to the protrusive spiral waves in a previously quiescent MCF-10A cell after the addition of rapamycin, which induces the recruitment of Inp54p-FKBP to the plasma membrane. Related to **Figure 4A**.

**Video S14.** Recruitment of GFP-FKBP-PFK (left) after the addition of rapamycin initiates cell spreading and F-actin wave activity (right, LifeAct-iRFP) in an MCF-10A M3 cell. Related to **Figure 4D**.

**Video S15.** An HL-60 neutrophil-like cell expressing CRY2PHR-mCherry-aldolase and LifeAct-miRFP703, before and after 488 nm light illumination. The HL-60 cell becomes highly motile and polarizes after aldolase recruitment to the plasma membrane. Related to **Figure 4G**.

**Video S16.** Video of two different MCF-10A M3 cells with 3 different focal planes showing the translocation of iRFP-FKBP-PFK (top) after addition of rapamycin induces cell spreading and also the recruitment of aldolase-GFP (bottom) to the plasma membrane. Related to **Figure 4K**.

